# Seasonal prevalence and characterization of Shiga toxin-producing *Escherichia coli* on pork carcasses at three steps of the harvest process at two commercial processing plants in the US

**DOI:** 10.1101/2020.07.15.205773

**Authors:** Ivan Nastasijevic, John W. Schmidt, Marija Boskovic, Milica Glisic, Norasak Kalchayanand, Steven D. Shackelford, Tommy L. Wheeler, Mohammad Koohmaraie, Joseph M. Bosilevac

## Abstract

Shiga toxin (*stx*) -producing *Escherichia coli* (STEC) are foodborne pathogens that have a significant impact on public health, with those possessing the attachment factor intimin (*eae*) referred to as enterohemorrhagic *E. coli* (EHEC) associated with life threatening illnesses. Cattle and beef are considered typical sources of STEC, but their presence in pork products is a growing concern. Therefore, carcasses (n=1536) at two U.S. pork processors were sampled once per season at three stages of harvest (post-stunning skins; post-scald carcasses; chilled carcasses) then examined using PCR for *stx* and *eae*, aerobic plate count (APC) and *Enterobacteriaceae* counts (EBC). Skins, post-scald, and chilled carcasses had prevalence of *stx* (85.3, 17.5, and 5.4%, respectively), with 82.3, 7.8, and 1.7% respectively, having *stx* and *eae* present. All *stx* positive samples were subjected to culture isolation that resulted in 368 STEC and 46 EHEC isolates. The most frequently identified STEC were serogroup O121, O8, and O91(63, 6.7, and 6.0% of total STEC, respectively). The most frequently isolated EHEC was serotype O157:H7 (63% of total EHEC). Results showed that scalding significantly reduced (*P* < 0.05) carcass APC and EBC by 3.00 and 2.50 log_10_ CFU/100 cm^2^ respectively. A seasonal effect was observed with STEC prevalence lower (*P* < 0.05) in winter. The data from this study shows significant (*P* < 0.05) reduction in the incidence of STEC (*stx*) from 85.3% to 5.4% and of EHEC (*stx*+*eae*) from 82.3% to 1.7% within slaughter-to-chilling continuum, respectively, and that potential EHEC can be confirmed present throughout using culture isolation.

**IMPORTANCE:** Seven serogroups of Shiga toxin-producing *Escherichia coli* (STEC) are responsible for most (>75%) cases of severe illnesses caused by STEC and are considered adulterants of beef. However, some STEC outbreaks have been attributed to pork products although the same *E. coli* are not considered adulterants in pork because little is known of their prevalence along the pork chain. The significance of the work presented here is that it identifies disease causing STEC, enterohemorrhagic *E. coli* (EHEC), demonstrating that these same organisms are a food safety hazard in pork as well as beef. The results show that most STEC isolated from pork are not likely to cause severe disease in humans and that processes used in pork harvest, such as scalding, offer a significant control point to reduce contamination. The results will assist the pork processing industry and regulatory agencies to optimize interventions to improve the safety of pork products.

## INTRODUCTION

Shiga toxin-producing *Escherichia coli* (STEC) are a potential food-borne pathogen that, after ingestion, can cause severe damage to the intestinal mucosa and, in some cases, other internal organs of the human host (1-3). Certain STEC possess adherence factors, the most commonly observed being intimin (*eae*), that contribute to the ability of the organism to cause enterohemorrhagic diseases such as hemorrhagic colitis which may develop into the life-threatening condition of hemolytic uremic syndrome (HUS). Since the early 1980s, *E. coli* O157:H7 has emerged as the enterohemorrhagic *E. coli* (EHEC) of the most significant public health relevance; not because of the incidence of the illness, which is much lower than that of other food-borne pathogens e.g. *Campylobacter* or *Salmonella*, but because of the severity of the symptoms, the low infectious dose and potential sequelae. Although the major source of STEC and EHEC are healthy ruminants, predominantly cattle, the increasing trend of foodborne outbreaks associated with *E. coli* O157:H7(O157-EHEC) and non-O157 EHEC that were reported over recent years, both in the USA and EU, were attributed to the consumption of pork (4-6).

In the USA, the annual testing of meat and meat products by the U.S Department of Agriculture (USDA) Food Safety and Inspection Service (FSIS) is designed to allow regular testing for product produced in domestic establishments, imported products, and raw ground beef in retail; the presence of O157*-*EHEC in samples of raw non-intact ground beef products and raw beef intended for raw non-intact products, including ground beef, raw ground beef components, and beef trimmings is carried out on a regular basis (7). The annual testing scheme also includes testing of raw pork meat for the presence of O157-EHEC, non-O157 EHEC and indicator organisms; 3800 samples of raw pork meat were tested in 2018, e.g. comminuted pork, intact pork cuts and non-intact pork cuts (7). In a recent report, of 1395 pork samples examined by FSIS for STEC, 309 (22%) screened positive for the presence of Shiga toxin genes (*stx*) and *eae*, but only 3 (0.2%) were confirmed by culture isolation (8). Unlike U.S. beef processors, U.S. pork processors do not conduct their own testing of products for *E. coli* O157:H7. At the moment in the EU, the only existing microbiological criterion for STEC in a food commodity is defined in the Regulation (EC) No. 209/2013 amending Regulation (EC) No. 2073/2005 as regards microbiological criteria for sprouts (9). The monitoring data on STEC in foods other than sprouts and in animals, originate from the reporting obligations of the EU Member States (10), which stipulates that Member States must investigate the presence of STEC at the ‘most appropriate stage’ of the food chain. Currently, Harmonized Epidemiological Indicators (HEI) at the EU level do not exist, allowing EU member states to carry out sampling, testing, data analysis and interpretation of results in a consistent manner.

In addition, the epidemiology and virulence factors of STEC and EHEC carried by on-farm pigs remain largely unknown. It is known healthy pigs are important reservoirs of STEC (11) and some isolated strains were reported as potential human pathogens (12, 13). Since certain outbreaks of STEC and EHEC were associated with pork consumption (6, 14-17), it is important to obtain additional scientific evidence on pathways of pork contamination by serogroups able to infect humans (18).

Thus, the aims of this study were: a) to determine the seasonal prevalence of STEC and EHEC as well as Aerobic plate count (APC) bacteria and *Enterobacteriaceae* counts (EBC) on pork carcasses at three different steps of harvest; b) to further characterize isolated STEC and EHEC strains; and c) to discuss the results obtained with their relevance to food safety and to propose the most effective control options for prevention/minimization of pork carcass contamination.

## RESULTS

### APC and EBC

Differences in the levels of APC and EBC of pork carcasses along the processing line at three points were observed between plants A and B (Table 1). During slaughter, the APC were higher (6.50 log_10_ CFU/100 cm^2^ in the plant A and 6.93 log_10_ CFU/100 cm^2^ in the plant B, respectively) on the carcass skin, while their numbers were significantly decreased (*P* < 0.05) following the scalding process (3.91 log_10_ CFU/100 cm^2^ in the plant A and 3.53 log_10_ CFU/100 cm^2^ in the plant B, respectively) and following final interventions when measured on chilled carcasses (2.48 log_10_ CFU/100 cm^2^ in the plant A and 2.22 log_10_ CFU/100 cm^2^ in the plant B, respectively). Carcass skin samples from the plants A and B-had EBC of 4.41 and 4.37 log_10_ CFU/100 cm^2^, respectively, while the carcasses showed significantly lower numbers of EBC after scalding (2.28 log_10_ CFU/100 cm^2^-plant A and 1.50 log_10_ CFU/100 cm^2^-plant B), and again in the chiller (0.88 log_10_ CFU/100 cm^2^-plant A and 0.49 log_10_ CFU/100 cm^2^-plant B) (*P* < 0.05).

**TABLE 1.** Aerobic Plate Counts (APC) and *Enterobacteriaceae* counts (EBC)^a^ on pork carcasses by sample site, processing plant, and season

| Season | Plant | APC (Log<sub>10</sub> CFU/100cm<sup>2</sup>) |  |  | EBC (Log<sub>10</sub> CFU/100cm<sup>2</sup>) |  |  |
| --- | --- | --- | --- | --- | --- | --- | --- |
|  |  | Skin<sup>c</sup> | Post-scald<sup>d</sup> | Final<sup>e</sup> | Skin | Post-scald | Final |
| - | A | 6.50b | 3.91a | 2.48a | 4.41a | 2.28a | 0.88a |
|  | B | 6.93a | 3.53b | 2.22b | 4.37a | 1.50b | 0.49b |
| Winter | - | 6.27y | 3.28x | 1.92y | 4.06y | 1.66y | 0.49y |
| Spring | - | 6.79x | 2.85z | 1.80y | 4.51x | 1.85x | 0.51y |
| Summer | - | 7.85w | 5.59w | 3.15w | 5.01w | 2.56w | 1.02w |
| Fall | - | 5.95z | 3.05y | 2.53x | 3.99z | 1.77xy | 0.73x |

<sup>a</sup>Values represent the mean Log<sub>10</sub> CFU/100cm<sup>2</sup> (n=768 by Plant and n=384 by Season), those
followed by the same letter within the column for plant or season are not different ( $P > 0.05$ ).
<sup>b</sup>Seasons: Winter = December-February, Spring = March-May, Summer = June-August, Fall = September-November.
<sup>c</sup>Skin of stunned exsanguinated pigs sampled along belly midline.
<sup>d</sup>Post-scald pre-evisceration pig carcasses sampled along midline from ham to breast, including fore shank and jowl. Carcass samples are not matched to other samples.
<sup>e</sup>Final = chilled finished pig carcasses, sampled along the split midline from ham collar to jowl and fore shank. Carcass samples are not matched to other samples.

Season significantly influenced (*P* < 0.05) skin contamination. Significantly higher APC and EBC were measured on carcass surfaces during summer (7.85 and 5.01 log_10_ CFU/cm^2^, respectively), followed by spring (6.79 and 4.51 log_10_ CFU/cm^2^) and winter (6.27 and 4.06 log_10_ CFU/cm^2^), while the lowest number of these bacteria were found during fall (5.95 and 3.99 log_10_ CFU/cm^2^). Although scalding significantly decreased number of these bacterial groups, seasonal variations remained significant (*P* < 0.05). After all interventions, carcasses in the chiller had the lowest numbers of APC and EBC recorded during winter (1.92 and 0.49 log_10_ CFU/cm^2^, respectively) and spring (1.80 and 0.51 log_10_ CFU/cm^2^) with no significant differences (*P* > 0.05) observed between these two seasons.

### PCR screening of pork carcasses for STEC (*stx*) and EHEC (*stx+eae*)

All samples were enriched then screened by PCR for Shiga toxin (*stx*) and intimin (*eae*) genes. The presence of *stx* was considered to indicate the presence of a STEC, while the concomitant presence of *eae* identified samples that potentially contained EHEC. Therefore, a sample that was PCR screened positive for *stx* and *eae* was included in both the potential STEC positive and the potential EHEC positive groups. In regard to STEC and EHEC screening of skins, post-scald pre-evisceration carcasses, and final carcasses, seasonal and plant differences were observed (Table 2).

**TABLE 2.** Prevalence^a^ of STEC^b^ and EHEC^c^ in samples collected from pork processing as determined by PCR^d^

| Season<sup>e</sup> | Plant | n | STEC Screen Positive |  |  | EHEC Screen Positive |  |  |
| --- | --- | --- | --- | --- | --- | --- | --- | --- |
|  |  |  | Skin<sup>f</sup> | Post-scald<sup>g</sup> | Final<sup>h</sup> | Skin | Post-scald | Final |
| - | - | 1536 | 85.3 | 17.5 | 5.4 | 82.3 | 7.8 | 1.7 |
| - | A | 768 | 81.3y<sup>f</sup> | 13.8y | 8.2x | 76.3y | 7.7x | 3.1x |
| - | B | 768 | 89.3x | 21.2x | 2.6y | 88.3x | 7.9x | 0.3y |
| Winter | - | 384 | 41.7r | 20.3q | 3.6qr | 29.7r | 9.6q | 0.8qr |
| Spring | - | 384 | 100.0q | 11.2r | 3.4r | 100.0q | 2.9r | 0.0r |
| Summer | - | 384 | 99.5q | 19.0q | 7.6q | 99.5q | 8.3q | 3.4q |
| Fall | - | 384 | 100.0q | 19.5q | 7.0qr | 100.0q | 10.4q | 2.6qr |
| Winter | A | 192 | 26.0c | 12.5gf | 5.2ih | 6.3de | 7.8cde | 1.6gh |
|  | B | 192 | 57.3b | 28.1c | 2.1i | 53.1b | 11.5c | 0.0h |
| Spring | A | 192 | 100.0a | 8.3gh | 5.2ih | 100.0a | 4.2efg | 0.0h |
|  | B | 192 | 100.0a | 14.1gef | 1.6i | 100.0a | 1.6fgh | 0.0h |
| Summer | A | 192 | 99.0a | 18.2def | 13.0gf | 99.0a | 8.3cd | 5.7de |
|  | B | 192 | 100.0a | 19.8de | 2.1i | 100.0a | 8.3cd | 1.0gh |
| Fall | A | 192 | 100.0a | 16.2ef | 9.4gh | 100.0a | 10.4c | 5.2dfe |
|  | B | 192 | 100.0a | 22.9dc | 4.7ih | 100.0a | 10.4c | 0.0h |

<sup>a</sup>Values represent percentages of each sample type in each category found positive.
<sup>b</sup>STEC are Shiga toxin-producing *E. coli* indicated by the presence of *stx1* and or *stx2* gene(s) in the sample.
<sup>c</sup>EHEC are enterohemorrhagic *E. coli* indicated by the presence of Shiga toxin (*stx*) and intimin (*eae*) genes in the sample.
<sup>d</sup>The screening PCR identified *stx1*, *stx2*, and *eae* genes in the enriched samples.
<sup>e</sup>Seasons: Winter = December-February, Spring = March-May, Summer = June-August, Fall = September-November.
<sup>f</sup>Skin of stunned exsanguinated pigs sampled along belly midline.
<sup>g</sup>Post-scald pre-evisceration pig carcasses sampled along midline from ham to breast, including fore shank and jowl. Carcass samples are not matched to other samples.
<sup>h</sup>Final = chilled finished pig carcasses, sampled along the split midline from ham collar to jowl and fore shank. Samples are not matched to other samples.
<sup>f</sup>Values within a group, STEC or EHEC, Plant (columns), Season (columns), or Plant x Season (columns and rows) followed by the same letter are not different ( $P > 0.05$ ).

Overall, skin samples were 85.3% positive for STEC, but with Plant A having a lower rate (*P* < 0.05) than Plant B. Seasonally, nearly 100% of skin samples were positive year-round for STEC except for the winter months when STEC prevalence was 41.7% (*P* < 0.05). During the winter, prevalence of STEC at Plant A was 26.0%, half that of Plant B (57.3%). This winter difference was responsible for all other differences observed on skins.

Following scalding and singeing but before any further processing, 17.5% of the pre-evisceration carcasses were STEC screen positive. Again, Plant A had a lower rate (13.8%) and was different (*P* < 0.05) from Plant B (21.2%). The seasonal effect observed on these carcasses was different however, from that of the incoming skins. While winter month skins screened lower for STEC, spring post-scald carcasses (11.2%) were lower (*P* < 0.05) than the other seasons (19-20%). The lowest post-scald carcass STEC screen rate was observed at Plant A in the spring (8.3%) while the highest was observed at Plant B in the winter (28.1%). Just 5.4% of the final carcasses in the chillers at Plants A and B combined screened positive for STEC, with Plant A having approximately a three-fold greater STEC prevalence (*P* < 0.05) than Plant B. Seasonally, summer final carcasses possessed the greatest number of STEC positives (7.6%), with the lowest (*P* < 0.05) number of STEC positives in the spring (3.4%). However, rates in the winter and fall, 3.6% and 7.0%, respectively, were not different (*P* > 0.05) than the summer and spring levels, respectively. The seasonally observed rates of STEC positive final carcasses at Plant A ranged from 5.2 to 13.0% while at Plant B they ranged from 1.6 to 4.7%.

Since potential EHEC screen positive samples represent a subset of all STEC screen positive samples, the prevalence of potential EHEC on skins and the carcasses was lower, however the plant and seasonal differences were generally maintained. Pork skins that screened positive for both *stx* and *eae* were 82.3%, Plant A (76.3%) and Plant B (88.3%) being different (*P* < 0.05); and winter skins (29.7%) less (*P* < 0.05) than the other seasons (99.5-100%). Nearly all skin samples were positive for both markers indicating presence of potential EHEC except for winter where only 6.3% of Plant A and 53.1% of Plant B skin samples screened positive for potential EHEC.

Post-scald carcasses screened positive for potential EHEC at 7.8% and were not different (*P* > 0.05) between the two plants (7.7 and 7.9%). There was a seasonal effect that followed the STEC positive screening with spring lower (2.9%; *P* < 0.05) than the three other seasons which were not different (*P* > 0.05) from one another and ranged from 8.3 to 10.4% potential EHEC screen positive.

The screened potential EHEC prevalence for final carcasses was very low at only 1.7%, but with significant differences (*P* < 0.05) between Plant A at 3.1% and Plant B at 0.3%. No final carcasses screened positive for EHEC in the spring months, whereas 3.4% of final carcasses did so in the summer months. This was the only seasonal effect observed amongst final carcasses. In a season by Plant analysis, Plant B only had final carcasses screen positive for potential EHEC in the summer at 1.0%, whereas Plant A potential EHEC screening identified 1.6% of final carcasses in the winter, which was less than the summer rate of 5.7% and significantly less (*P* < 0.05) than the fall rate (5.2%).

### Isolation of STEC and EHEC from pork processing samples

The presence of an EHEC exclusive of a STEC could only be confirmed by culture isolation, as the samples could have been co-contaminated by a STEC (possessing an *stx* gene) and an atypical enteropathogenic *E. coli* (EPEC; possessing an *eae* gene). Therefore, all *stx* PCR screen positive samples were subjected to culture confirmation. In total, 405 samples were culture confirmed. Three hundred sixty (360) of the samples yielded 368 different STEC isolates (Table 3) while 46 samples yielded 46 EHEC isolates (Table 4). One sample was culture confirmed to harbor a STEC and an EHEC. Most isolates were found in samples collected in the spring and summer months, 120 and 135, respectively. Whereas, only 67 winter samples and 92 fall samples were culture confirmed. O121 was the most common STEC on skin and post-scald carcasses and O157 was the most common EHEC.

**TABLE 3.** Summary^a^ of STEC^b^ strains (n=368) isolated from pork processing plants by sample type, season^c^, and processing plant

|  |  | STEC Serogroup |  |  |  |  |  |  |  |  |  |  |  |  |  |  |  |  |  |
| --- | --- | --- | --- | --- | --- | --- | --- | --- | --- | --- | --- | --- | --- | --- | --- | --- | --- | --- | --- |
|  | Season | Plant | O2 | O5 | O8 | O20 | O32 | O55 | O74 | O86 | O91 | O103 | O110 | O112 | O121 | O139 | O141 | O146 | ONT<sup>g</sup> |
| Skin<sup>d</sup> |  |  |  |  |  |  |  |  |  |  |  |  |  |  |  |  |  |  |  |
|  | Winter | A | 1 |  | 15 | 7 |  |  |  |  |  |  | 1 |  | 15 | 1 |  |  | 2 |
|  |  | B |  |  |  |  | 1 |  |  |  | 10 |  |  |  | 1 | 4 |  | 1 | 4 |
|  | Spring | A |  |  | 1 |  |  |  |  |  |  |  |  | 1 | 26 |  |  |  | 5 |
|  |  | B |  |  | 4 |  |  | 7 |  |  | 5 |  |  |  | 50 | 5 | 2 |  | 13 |
|  | Summer | A |  |  | 1 | 1 |  |  | 1 |  | 3 |  |  |  | 17 | 1 |  |  | 1 |
|  |  | B |  | 1 | 1 | 1 |  |  |  |  | 3 | 1 |  |  | 68 |  |  | 2 | 7 |
|  | Fall | A |  |  |  | 1 |  |  |  |  |  |  |  |  | 6 |  | 1 |  | 1 |
|  |  | B |  |  | 1 |  |  |  |  |  |  |  |  |  | 42 | 3 |  |  | 3 |
| Post-scald carcass<sup>e</sup> |  |  |  |  |  |  |  |  |  |  |  |  |  |  |  |  |  |  |  |
|  | Summer | A |  |  | 1 |  |  |  |  |  |  |  |  |  | 1 |  |  |  |  |
|  |  | B |  |  | 1 |  |  |  |  | 1 |  |  |  |  | 1 |  |  |  | 3 |
|  | Fall | A |  |  |  |  |  |  |  | 1 |  |  |  |  | 2 |  |  |  |  |
|  |  | B |  |  |  |  |  |  |  |  | 1 |  |  |  | 2 |  |  |  |  |
| Final Carcass<sup>f</sup> |  |  |  |  |  |  |  |  |  |  |  |  |  |  |  |  |  |  |  |
|  | Summer | A |  |  |  |  |  |  |  |  |  |  |  |  | 1 | 1 |  |  | 3 |
|  |  | B |  |  |  |  |  |  |  |  |  |  |  |  |  |  |  |  |  |
| Total |  |  | 1 | 1 | 25 | 10 | 1 | 7 | 1 | 2 | 22 | 1 | 1 | 1 | 232 | 15 | 3 | 3 | 42 |

<sup>a</sup>Values represent the number of isolates recovered from samples within each category.
<sup>b</sup>STEC are Shiga toxin-producing *E. coli* lacking intimin (*eae*) gene.
<sup>c</sup>Seasons: Winter = December-February, Spring = March-May, Summer = June-August, Fall = September-November.
<sup>d</sup>Skin of stunned exsanguinated pigs sampled along belly midline.
<sup>e</sup>Post-scald pre-evisceration pig carcasses sampled along midline from ham to breast, including fore shank and jowl. Carcasses are not matched to other samples.
<sup>f</sup>Final = chilled finished pig carcasses, along the split midline from ham collar to jowl and fore shank. Carcasses are not matched to other samples.
<sup>g</sup>ONT=serogroup was not typable using limited antisera sets available.

**TABLE 4.** Summary^a^ of EHEC^b^ (n=46) isolated from pork processing plants by season^c^, and processing plant

| Season | Plant | EHEC serogroup |  |  |  |  |  |
| --- | --- | --- | --- | --- | --- | --- | --- |
|  |  | O8 | O26 | O103 | O121 | O157 | ONT<sup>d</sup> |
| Winter | A |  |  | 2 |  |  |  |
|  | B |  |  |  |  |  | 2 |
| Spring | A |  |  |  |  |  | 1 |
|  | B |  |  |  |  |  |  |
| Summer | A |  | 2<sup>e</sup> |  |  |  |  |
|  | B |  |  |  | 2 | 8 | 1 |
| Fall | A | 1 |  |  | 3 | 15 | 1<sup>e</sup> |
|  | B |  | 1 |  | 1 | 6<sup>e</sup> |  |
| Total |  | 1 | 3 | 2 | 6 | 29 | 5 |

<sup>a</sup>Values represent number of EHEC isolates of the given serogroup recovered from samples that screened positive for Shiga toxin genes by PCR.
<sup>b</sup>EHEC are enterohemorrhagic *E. coli* possessing Shiga toxin (*stx*) and intimin (*eae*) genes.
<sup>c</sup>Seasons: Winter = December-February, Spring = March-May, Summer = June-August, Fall = September-November.
<sup>d</sup>ONT = serogroup was not typable using limited antisera sets available.
<sup>e</sup>All isolates were recovered from Pork Skin swab samples except the 2 EHEC-O26 (Plant A, Summer) that were recovered from final pork carcasses; 1 EHEC-ONT (Plant A, Fall) recovered from a pre-intervention carcass; and 1 EHEC-O157 (Plant B, Fall) recovered from a pre-intervention carcass.

As suggested by the PCR screening results, samples collected from skins yielded the most STEC and EHEC isolates (Tables 3 and 4). Plant B had about twice as many skin samples culture confirmed with a STEC (n=240) compared to Plant A (n=109), but both plants had similar number of skin samples culture confirmed an EHEC (25 and 21 for Plants A and B, respectively). Samples collected in the spring, and winter months only yielded 4 and 1 EHEC, respectively, with the bulk of the isolated EHEC being found in the summer and fall (Table 4).

Nearly two-thirds (64.4%) of the STEC isolated from skins were STEC-O121. STEC with non-typeable serogroups were second most common (10.5%). These two groups of STEC were the only ones found at both plants every season. Other STEC identified at both plants and/or during more than one season were STEC-O8, O91, O139, and O20 (Table 3).The most common EHEC isolated from skins was EHEC-O157:H7, which made up 63.0% of the EHEC isolates from skins. EHEC-O157:H7 was found at Plant B in the summer and both plants in the fall. The next most common EHEC isolated from skin samples was EHEC-O121. It too was isolated in a similar pattern as that of EHEC-O157:H7. Other EHEC isolated from skins were O8, O26, O103 and O-non-typeable (Table 4).

Post-scald pre-evisceration carcasses were found 17.5% PCR screen positive for a STEC and culture confirmed at a rate of 0.9%, while 1.7% PCR screened positive for a potential EHEC but only 0.1% culture confirmed for an EHEC. All isolates from post-scald carcasses were only recovered from samples collected in the summer and fall months. These were the seasons with some of the highest PCR screen positive rates. A third fewer STEC were found at Plant A in the summer than at Plant B. However, STEC-O8 and STEC-O121 were present at both plants in the summer. Similar numbers of STEC were found at each plant in the fall, again with STEC-O121 being most common. One EHEC was isolated from the post-scald carcasses at each plant in the fall. These isolates were an EHEC-O157:H7 at Plant B and an EHEC-ONT at Plant A.

Final carcasses also only had 5 STEC isolated, STEC-O121, O139, and 3 ONT recovered from Plant A during the summer. Only 2 EHEC-O26 were culture confirmed from final carcasses, similarly from Plant A during the summer. No isolates were recovered from final carcasses at Plant B, nor during any other season. The recovery of isolates agrees with the PCR screening results being highest for Plant A in the summer at 13.0 and 5.7% for STEC and potential EHEC, respectively.

### Characterization of STEC isolates

Of the 367 STEC isolated, 6 were recovered from post-scald carcasses and 1 from a final carcass, while the remaining 360 isolates were found on pre-scald carcass skins. STEC-O121 made up 63% of the isolates (Table 5). Eighteen variations were observed based on the presence of the different virulence factors examined. Seven of the genotypes were unique isolates, whereas multiple isolates of similar genotypes numbered in groups of 2 to 163. In the case of 6 genotypes the identical isolates were found across plants and seasons. However, one genotype represented by 163 isolates were all recovered from skin samples at Plant A during the spring. All but 7 of the STEC-O121 isolates (6 from skin, 1 from post-scald carcass) possessed Shiga toxin 2 subtype e (*stx*_*2e*_). Two isolates carried an *stx*_*1a*_ allele in addition to the *stx*_*2e*_ allele. Only 5 STEC-O121 possessed what appeared to be incomplete pO157 plasmids. All five carried *katP*, while two also possessed *etpD*, with one of those also having *espP* (Table 5). Most of the STEC-O121 carried an allele of *eastA*, and a small number also possessed iron acquisition genes. Two STEC-O121 possessed the adherence factor *saa*, these were found at Plant B in the fall and Plant A in the winter.

**TABLE 5.**
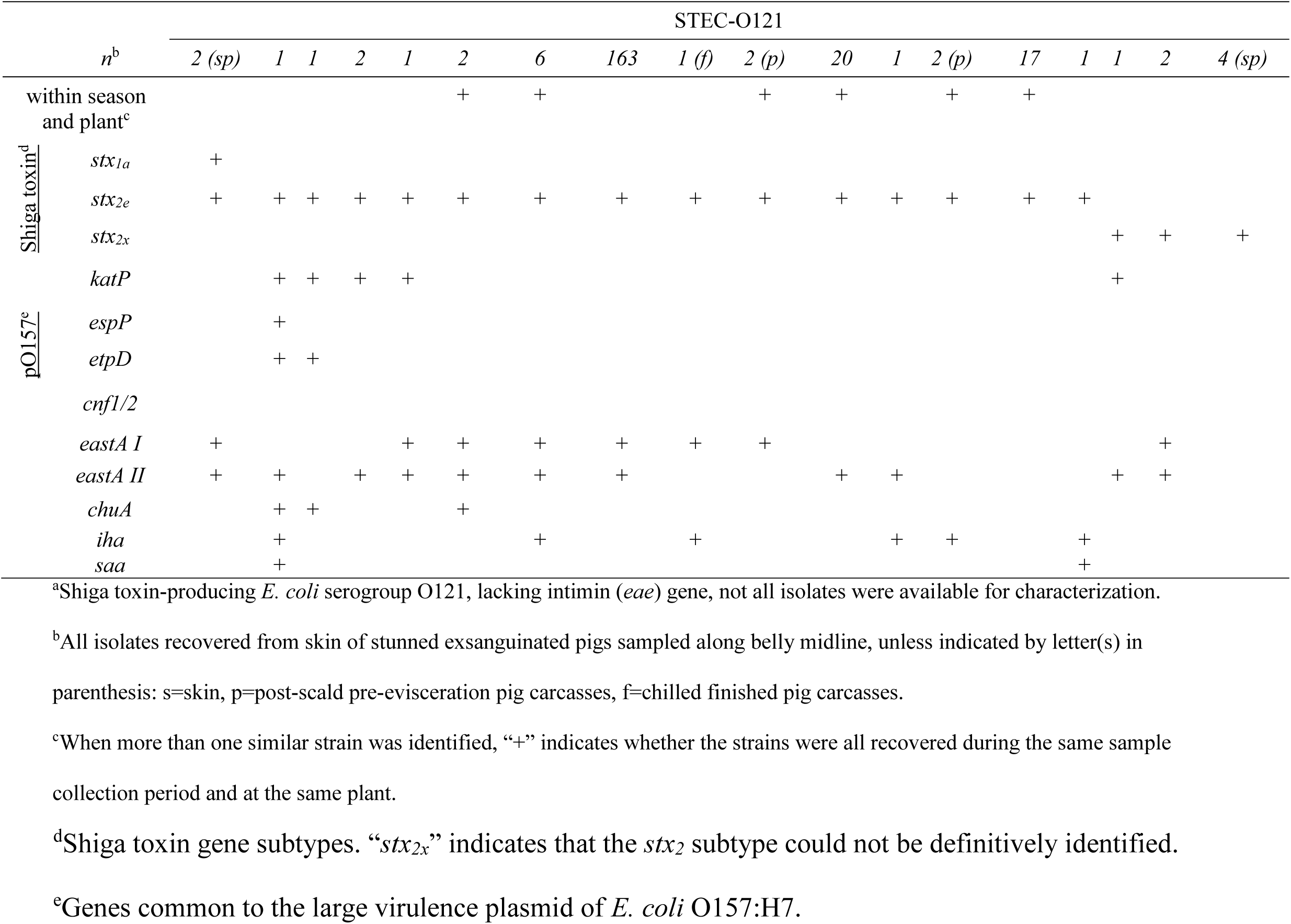
Characterization of STEC-O121(n=229)^a^ isolated from pork processing

The remaining STEC isolates (n=134) were of 15 serogroups and a large group (n=41) of non-identified serogroups (this due to our limited serotyping anti-sera). The identified serogroups included O2, O5, O8, O20, O32, O55, O74, O86, O91, O103 (an intimin lacking STEC), O110, O112, O139, O141, and O146. These STEC non-O121 (Table 6) also predominantly had *stx*_*2e*_. Shiga toxin subtype 1a (*stx*_*1a*_) was the lone Shiga toxin in 21 isolates of serogroups O20, O32, O91, O110, O112, and ONT. Shiga toxin subtypes 2a (*stx*_*2a*_) and 2c (*stx*_*2c*_) were uncommon, observed in only 2 isolates, a STEC-O8 and a STEC ONT, respectively. Six isolates had *stx*_*2*_ of non-identifiable subtypes. In most cases *stx* occurred as a single allele except for a STEC-O8 possessing *stx*_*2e*_ and *stx*_*2a*_, a STEC-O32 with *stx*_*1a*_ and *stx*_*2x*_, and STEC-ONTs that possessed combinations of *stx*_*1a*_ with *stx*_*2x*_, *stx*_*2c*_ with *stx*_*2x*_, and *stx*_*1a*_ with *stx*_*2e*_.

**TABLE 6.**
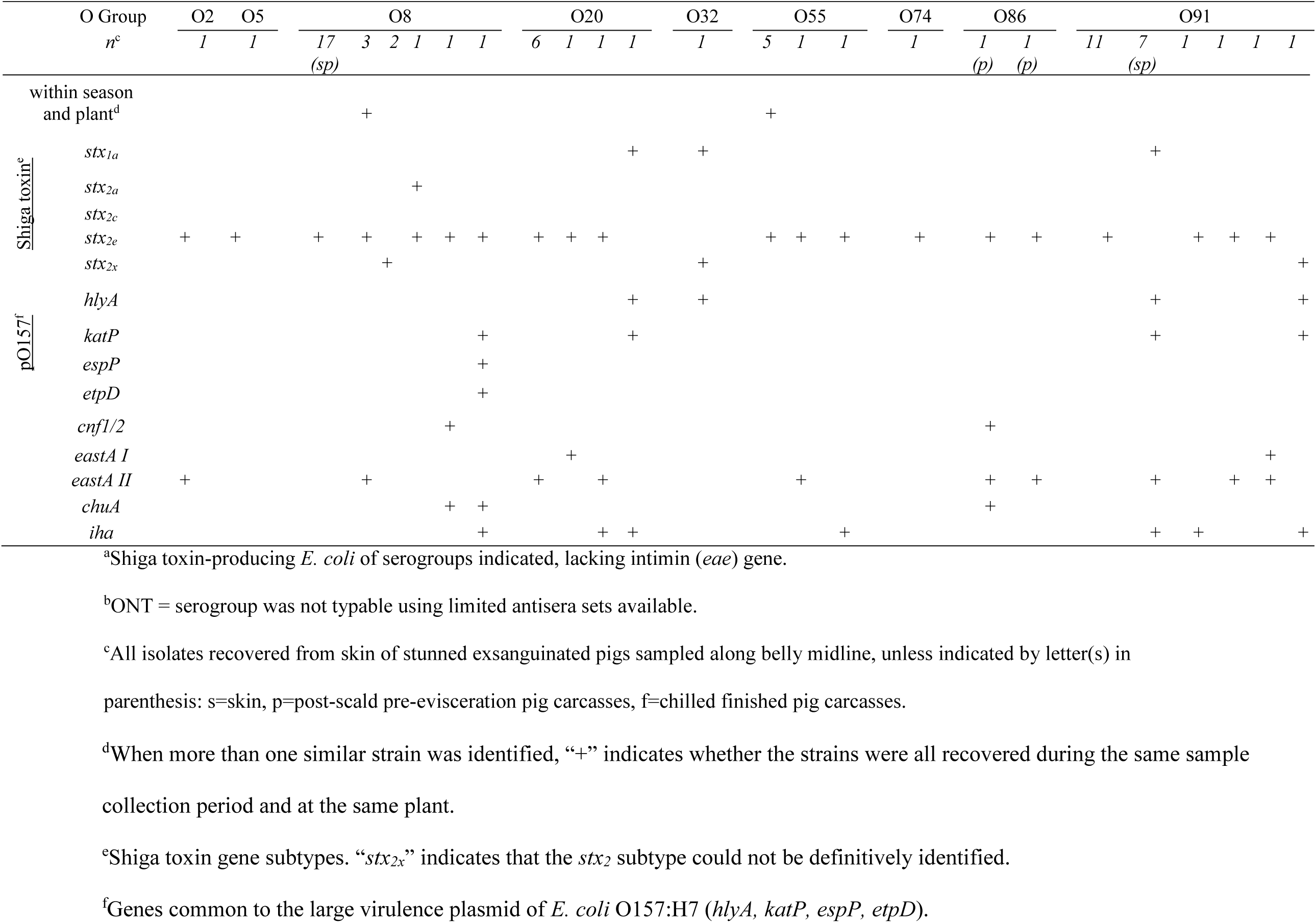
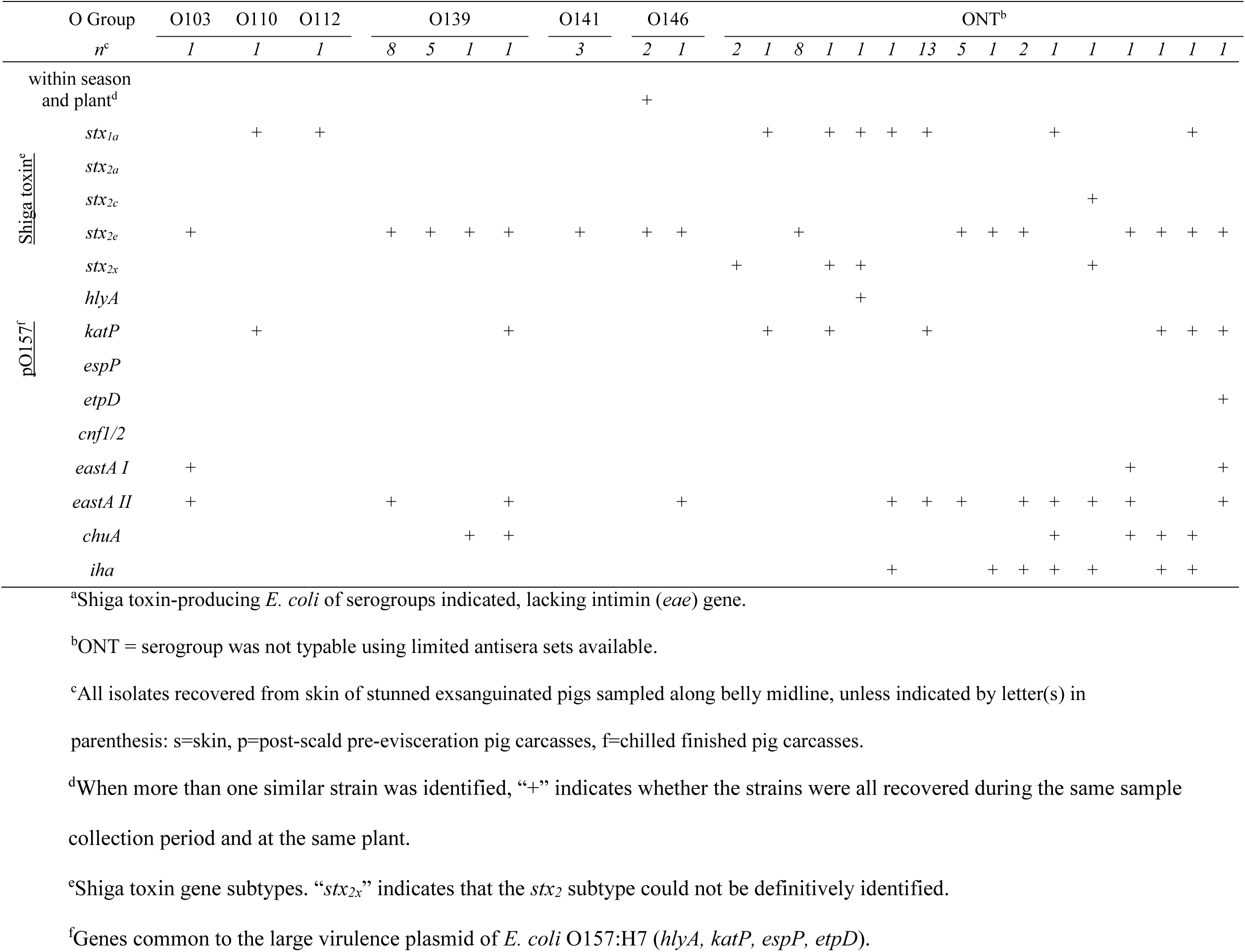
Characterization of STEC^a^ (non-O121) isolated from pork processing

Incomplete variations of the pO157 plasmid were observed in multiple isolates. Eight STEC-O91 possessed the pO157 markers *hlyA* and *katP*, and these were the two most common of the pO157 markers identified in the STEC isolates (30 had *katP* and 11 had *hlyA*). One STEC-O8 had three pO157 markers present (*katP, espP*, and *etpD*) and represented the most complete pO157 plasmid within the non-O121 STEC isolates. In regard to other virulence factors, 2 isolates possessed cytotoxic necrotizing factor (*cnf*), a STEC-O8 and a STEC-O86. Multiple strains had alleles of *eastA*, while iron acquisition genes *iha* and *chuA* were observed in isolates of STEC-O8, -O20, -O55, -O86, -O91, and -O139. Fourteen of the STEC-ONT lacked these additional factors, while the rest possessed 2 or more of them.

### Characterization of EHEC isolates

The EHEC isolates divided into *E. coli* O157:H7 (n=29; Table 7) and non-O157 EHEC (n=17; Table 8). The 29 *E. coli* O157:H7, when compared for Shiga toxin types, *nle* effectors, composition of the pO157 plasmid, and other toxin, adherence, and iron utilization genes, all impacting virulence, resulted in 12 different genotypes (Table 7).

**TABLE 7.**
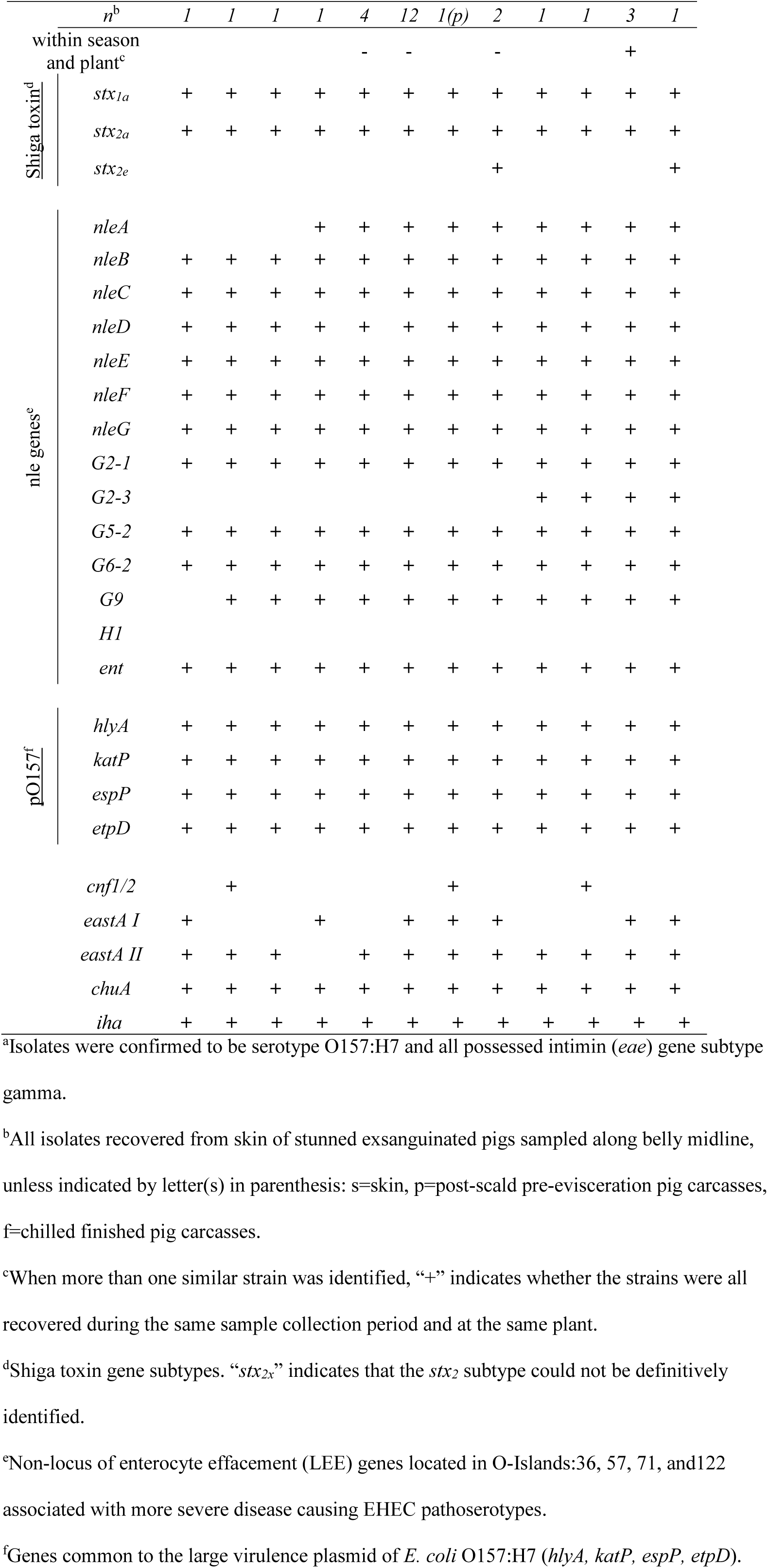
Characterization of EHEC O157:H7 isolates^a^

|  | *n*<sup>b</sup> | *1* | *1* | *1* | *1* | *4* | *12* | *1(p)* | *2* | *1* | *1* | *3* | *1* |
| --- | --- | --- | --- | --- | --- | --- | --- | --- | --- | --- | --- | --- | --- |
| within season<br>and plant<sup>c</sup> |  |  |  |  |  | - | - |  | - |  |  | + |  |
| <u>Shiga toxin<sup>d</sup></u> | *stx1a* | + | + | + | + | + | + | + | + | + | + | + | + |
|  | *stx2a* | + | + | + | + | + | + | + | + | + | + | + | + |
|  | *stx2e* |  |  |  |  |  |  |  | + |  |  |  | + |
| <u>nle genes<sup>e</sup></u> | *nleA* |  |  |  | + | + | + | + | + | + | + | + | + |
|  | *nleB* | + | + | + | + | + | + | + | + | + | + | + | + |
|  | *nleC* | + | + | + | + | + | + | + | + | + | + | + | + |
|  | *nleD* | + | + | + | + | + | + | + | + | + | + | + | + |
|  | *nleE* | + | + | + | + | + | + | + | + | + | + | + | + |
|  | *nleF* | + | + | + | + | + | + | + | + | + | + | + | + |
|  | *nleG* | + | + | + | + | + | + | + | + | + | + | + | + |
|  | *G2-1* | + | + | + | + | + | + | + | + | + | + | + | + |
|  | *G2-3* |  |  |  |  |  |  |  |  | + | + | + | + |
|  | *G5-2* | + | + | + | + | + | + | + | + | + | + | + | + |
|  | *G6-2* | + | + | + | + | + | + | + | + | + | + | + | + |
|  | *G9* |  | + | + | + | + | + | + | + | + | + | + | + |
|  | *H1* |  |  |  |  |  |  |  |  |  |  |  |  |
|  | *ent* | + | + | + | + | + | + | + | + | + | + | + | + |
| <u>pO157<sup>f</sup></u> | *hlyA* | + | + | + | + | + | + | + | + | + | + | + | + |
|  | *katP* | + | + | + | + | + | + | + | + | + | + | + | + |
|  | *espP* | + | + | + | + | + | + | + | + | + | + | + | + |
|  | *etpD* | + | + | + | + | + | + | + | + | + | + | + | + |
|  | *cnf1/2* |  | + |  |  |  |  | + |  |  | + |  |  |
|  | *eastA I* | + |  |  | + |  | + | + | + |  |  | + | + |
|  | *eastA II* | + | + | + |  | + | + | + | + | + | + | + | + |
|  | *chuA* | + | + | + | + | + | + | + | + | + | + | + | + |
|  | *iha* | + | + | + | + | + | + | + | + | + | + | + | + |

<sup>a</sup>Isolates were confirmed to be serotype O157:H7 and all possessed intimin (*eae*) gene subtype
gamma.
<sup>b</sup>All isolates recovered from skin of stunned exsanguinated pigs sampled along belly midline,
unless indicated by letter(s) in parenthesis: s=skin, p=post-scald pre-evisceration pig carcasses,
f=chilled finished pig carcasses.
<sup>c</sup>When more than one similar strain was identified, “+” indicates whether the strains were all
recovered during the same sample collection period and at the same plant.
<sup>d</sup>Shiga toxin gene subtypes. “*stx<sub>2x</sub>*” indicates that the *stx<sub>2</sub>* subtype could not be definitively
identified.
<sup>e</sup>Non-locus of enterocyte effacement (LEE) genes located in O-Islands:36, 57, 71, and 122

**TABLE 8.**
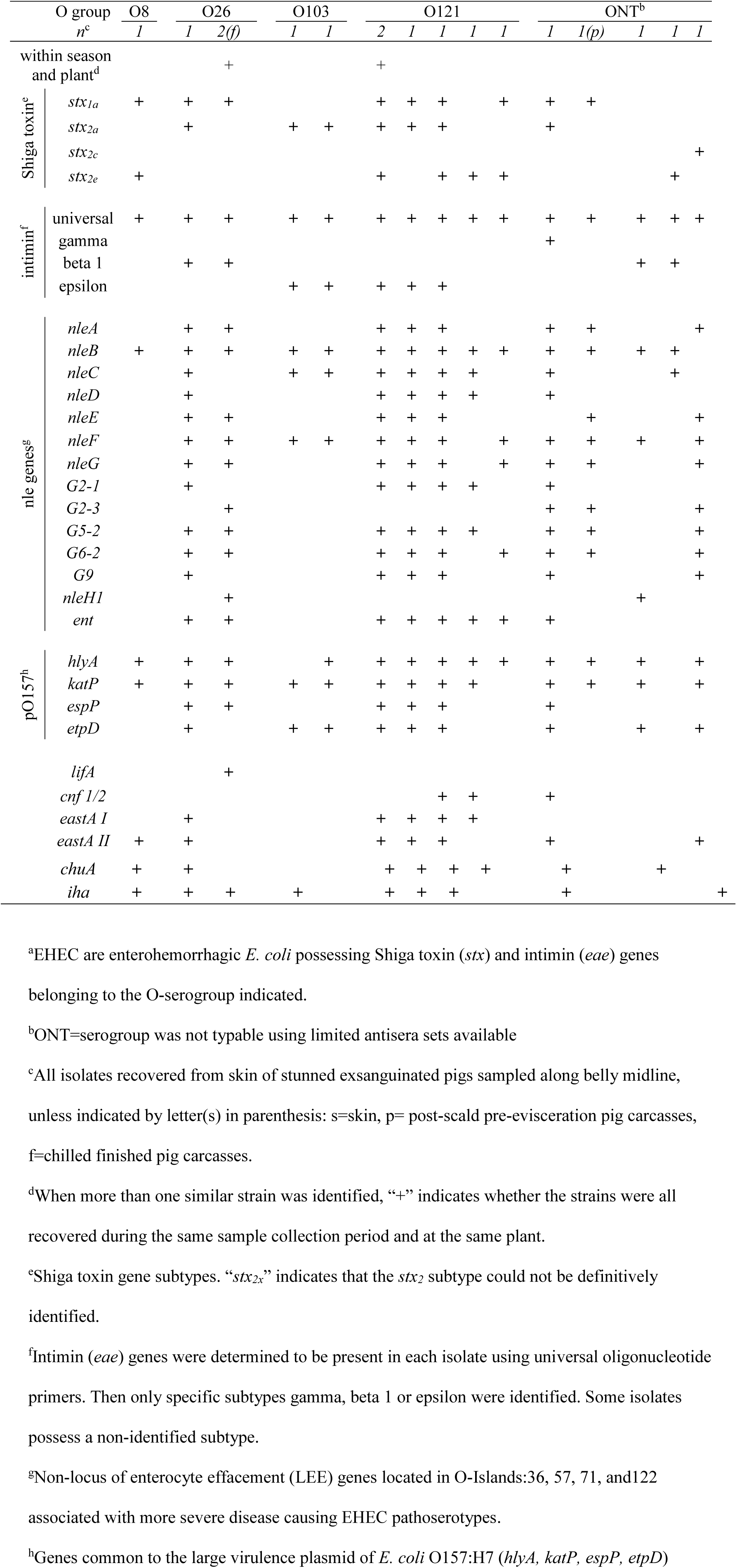
Characterization of non-O157 EHEC^a^ isolates

Twelve of the 29 *E. coli* O157:H7 isolates possessed identical patterns of the genes and were found across seasons and between the two plants. All the *E. coli* O157:H7 possessed *stx*_*1*_ and *stx*_*2a*_, but 3 isolates also carried the *stx*_*2e*_ allele (Table 7). All *E. coli* O157:H7 isolates appeared to possess an intact pO157 plasmid as evidenced by the presence of the *hylA, katP, espP* and *etpD* genes which are spaced around the plasmid. The iron utilization genes *chuA* and *iha* were also present in all of the *E. coli* O157:H7 isolates. The primary differences between the *E. coli* O157:H7 strains involved differences in the presence of the *nle* genes *nleA, nleG2-3*, and *nleG9* as well as cytotoxic necrotizing factor (present in 3) and *E. coli* heat stable enterotoxin 1. Non-O157 EHEC (n=17) were of 4 identifiable serogroups (O8, O26, O103 and O121) with 5 isolates having a non-typeable serogroup (Table 8). The non-O157 EHEC divided into fifteen groups based on genetic composition. These EHEC possessed different complements of Shiga toxin alleles, *stx*_*1a*_, *stx*_*2a*_, *stx*_*2c*_ and *stx*_*2e*_. Three of the most frequent non-O157 STEC serogroups recognized by the CDC (1) and FSIS(19) were identified (O26, O103 and O121), each possessing the expected *eae* subtypes of β1 and ε, however 2 of the EHEC-O121s had an *eae* gene that could not be subtyped using our primer sets suggesting it may be something other than *eae*-ε. Intimin-γ was observed in one of the EHEC-ONT. This isolate may be an EHEC-O145 that lacks the chromosomal region our serogrouping PCR identifies. This strain did not appear to have rfbO157, or flicH7 on PCR and was a sorbitol fermenter (data not shown) suggesting it is not likely an *E. coli* O157:H7.

Variable numbers of *nle* genes were observed in the EHEC isolates with the EHEC-O8 and 2 of the EHEC-ONT possessing only 1 to 3 of the effectors (Table 8). The 2 EHEC-O103 lacked many of the *nle* genes in comparison to the EHEC-O26s. Two of the EHEC-O121 and one of the EHEC-ONT possessed nearly all of the *nle* genes. Intact and partial pO157 plasmids were identified in the non-O157 EHEC. An EHEC-O26, 4 -O121, and an -ONT all appeared to possess a complete plasmid, while other isolates had incomplete versions. One EHEC-ONT lacked all markers for the pO157 plasmid. In regard to other factors, the *lifA* gene was only present in one EHEC-O26 found at Plant A during the summer. Cytotoxic necrotizing factor, and *E. coli* heat stable enterotoxin, and iron acquisition factors (*iha* and *chuA*) were variably present in all but four of the non-O157 EHEC isolated from pork carcasses.

## DISCUSSION

The present study identifies STEC and potential EHEC on skins of pre-scald pork carcasses in two U.S. commercial plants. Contamination of pigs with pathogenic EHEC-O157 and non-O157 may have occurred at farms (feed, water, manure), during transport, or lairage. Available data shows that some EHEC-O157 strains may persist for more than two years in farm environment (20). In addition, tonsils of some pigs have been reported to be colonized by significant levels of *E. coli* O157:H7 (21). The significantly higher (*P* < 0.05) STEC and EHEC prevalence on pre-scald carcasses sampled at Plant B could be due to higher contamination at any mentioned steps prior to slaughter, or potentially the “all in-all out” method of pork production where each farm empties a full facility for slaughter. However, determination of the source of this contamination was not the aim of the present study.

The results obtained in our study showed very high prevalence of the *stx* gene(s) indicating a STEC (85.3%) and the *stx* and *eae* genes indicating a potential EHEC (82.3%) on skin of pigs at slaughter. Nevertheless, significant decrease in prevalence of these genetic markers was observed after scalding in present study. Other authors reported the effectiveness of the scalding stage on decreases of *E. coli* and coliform counts on pork carcasses (22, 23). This important step is usually a Critical Control Point (CCP) within a risk-based food safety management system (Hazard Analysis and Critical Control Points/HACCP) and reduces both, the bacterial numbers and prevalence of pathogens (22).

APC bacteria are generally used to assess the hygiene of meat processing (24) and EBC are also used as indicators of fecal contamination (25, 26). The results of the present study showed that scalding is effective in reducing bacterial contamination on the carcass. Furthermore, our results are in the line with previous reports showing that scalding (59-62 °C) of pork carcasses resulted in reduction of APC (22, 27, 28). In other experiments scalding reduced APC and EBC by 3.1–3.8, and 1.7–3.3 log_10_ CFU cm^−2^, respectively (22, 27) which is similar to results found here (up to 3.4 log_10_ CFU 100cm^−2^ and 2.87 log_10_ CFU 100cm^−2^).

Unfortunately, epidemiological data on STEC prevalence in different regions and studies are not always comparable due to differences in study designs, different sampling and methods applied for detection and isolation, as well as season in which the study was performed (11, 18, 29). In Italy, Ercoli et al. (11), reported STEC prevalence of 13.8% on pork carcasses before chilling, while in Belgium prevalence of this pathogen was 12.8% on carcasses after cutting, before chilling (30). In the present study the prevalence of STEC after scalding ranged between 13.8% (Plant A) and 21.2% (Plant B). Moreover, the data from the present study also showed significant (*P* < 0.05) reduction in the incidence of STEC, indicated by *stx* gene(s), from 85.3% to 5.4% and of EHEC, indicated by *stx* and *eae* genes, from 82.3% to 1.7% within slaughter-to-chilling continuum, respectively. Colello et al. (29) found that 4.08% of pork carcasses sampled were *stx* positive in a study carried out in Argentina. Similar prevalence of STEC as in the present study (5.4%) was also found in carcasses after cooling in a Canadian study (4.8%) (31).

Since the complete elimination of pathogen(s) is not possible, chilling as an Standard Operating Procedure (SOP) has the objective, in general, to reduce carcass surface temperature preventing and slowing their growth (32, 33). In the present experiment, significant differences (*P* < 0.05) in carcass APC and EBC counts after chilling were observed between the two plants. These findings may be attributed to differences in chilling systems used by the plants. Although the incoming microorganism load on skins was higher at the beginning of harvest, at the end a lower level of APC and EBC and lower incidence of STEC was found in Plant B (2.22 and 0.49 log_10_ CFU/100cm^2^, 0.3%, respectively) where blast chilling was used, compared to conventional chilling in Plant A (2.48 and 0.88 log_10_ CFU/100cm^2^, 3.1%, respectively). Blast-chilling compared to conventional chilling lowers the carcass temperature at a rapid rate resulting in the arrest of contaminate growth when the population is smaller. In addition, blast chilling may provoke cold shock, especially in particularly sensitive Gram-negative microorganisms including *E. coli* and *Enterobacteriaceae*. Whereas, with conventional chilling, microorganisms may have opportunity to adapt to the low temperatures and avoid cold shock (34). However, the final carcasses that were sampled were not linked to the post-scald carcasses, and were in fact from lots of hogs harvested the previous days. The average reduction of APC from post-scald to final carcasses was not different (*P* > 0.05) between the two plants, while the reduction of EBC between these two points was significantly greater (*P* < 0.05) at Plant A (data not shown). Therefore, the significantly different microbial counts observed on carcasses in the chiller was likely a combination of the result of interventions applied as carcasses entered the chiller and the chilling process.

A lactic acid treatment following the final carcass water wash was applied as carcasses entered the chiller. It is well known that the combination of water and lactic acid treatment provide the greatest microbial reduction without large negative effects on quality attributes of pork meat (35, 36). As mentioned, in the present study carcasses in both plants were treated with 2% lactic acid (ambient temperature water, 10-30 s), before the cooling step. If the initial counts are higher, as in the present study, the effect of lactic acid decontamination treatment is more evident (36). Ba et al. (37) observed that significantly higher reductions in all bacterial species on the pork carcasses were achieved when sprayed with 4% lactic acid. Kalchayanand et al. (38) reported significant decrease of STEC O26, O45, O103, O111, O121, O145, O157 in inoculated fresh beef after lactic acid treatment.

Results regarding seasonal effect observed in the present study should be interpreted with caution because the visits to the plants were only carried out on two consecutive days during each period. It was observed that there were significant increases (*P* < 0.05) in APC and EBC during summer and spring compared to winter and fall. However, STEC prevalence indicated by *stx* genes on skin of pigs at harvest was high (99-100%) and did not differ between spring, summer and fall (*P* > 0.05). Only during winter was significantly lower prevalence (*P* < 0.05) of this pathogen indicator (*stx*) found compared to other seasons. Essendoubi et al. (26) also found higher prevalence of STEC on beef carcasses during warmer months (from June to November), while Dawson et al. (39) reported higher *E. coli* O157:H7 colonization in cattle during warmer months compared to cooler times of the year in various cattle production systems. One possible explanation may be that animals are dirtier during summer months due to soil and fecal contamination (33, 40, 41). In contrast, Cha et al. (42) reported higher STEC prevalence in pigs during fall and winter months (36.16% and 19.72%, respectively) suggested that low temperatures may contribute to increased stress in pigs leading to lower immunity and increased susceptibility to new STEC infections. The seasonal variations require further investigation as U.S. pigs are finished in doors in temperature controlled facilities and not directly exposed to colder temperatures in winter.

EHEC are important pathogens of public health significance because these isolates possess not only *stx*_*1*_ and/or the *stx*_*2*_ but also *eae*, the gene for the adherence factor intimin. Intimin, an integral outer membrane protein, is required for adherence and bacterial adhesion to enterocytes inducing a characteristic histopathological attaching and effacing (A/E) lesion and has been considered as a risk factor for disease in humans (29, 43). Although the presence of *eae* gene is an aggravating factor, this virulence factor was not always essential for severe illness suggesting that there might be also alternative mechanism of attachment (3). One such additional adherence factor we observed in a small number of STEC was *saa*, the STEC autoagglutinating adhesin. The *saa* gene had been identified in STEC isolated from humans with HUS or diarrhea (44).

Unlike strains that produce *stx*_*1*_, *stx*_*2*_-producing strains are often associated with HUS (45, 46). In the present study the strains possessing *stx*_*2*_ accounted for 88.74% of the total STEC isolates and 59.58% of all isolates (data not shown). While most *stx*_*2*_ genes were subtype *2e*, there were those that possessed *stx*_*2a*_ and *stx*_*2c*_, both major subtypes produced by *E. coli* strains associated with HUS (46). Strains that produce *stx*_*2e*_ do not consistently provoke foodborne illness in humans (47), but other data has confirmed isolation of *stx*_*2e*_-associated STEC from a HUS patient (48).With the exception of 8 STEC-O121 that had an unidentified *stx*_*2*_ subtype, the remaining STEC-O121 only possessed *stx*_*2e*_. STEC containing subtype *stx*_*2e*_ are typical swine-adapted STEC and present the most frequently reported Shiga toxin subtype from pigs (42, 49). This subtype is responsible for porcine edema disease in pigs (47) and consequently economic losses in production (13, 29).

EHEC serogroups isolated in the present study included O26 (3), O103 (2), O121 (5), and O157 (29).The USDA FSIS has declared the so called “big six” non-O157 serogroups (O26, O45, O103, O111, O121, O145) as adulterants in beef (19). These serotypes present public health burden because they are linked to a significant number of Hemorrhagic Colitis (HC) and Hemolytic Uremic Syndrome (HUS) cases (1, 50, 51). European Food Safety Authority (3) has made a similar declaration for serogroups with a high pathogenicity potential (O157, O26, O103, O145, O111, O145). Therefore, in present study the STEC serogroups of public health importance that were isolated were O157 and O103 (3) and O157, O26, O103, O121 (19).

Most of the EHEC isolates found in present study were O157:H7 (29) and were isolated from both plants during summer and fall. Serotype O157:H7 causes the most severe clinical symptoms. Although pork is not a common vehicle of EHEC-O157, some outbreaks of this infection in U.S., Canada (15-17, 52), and Italy (53) have been linked to consumption of roasted pork meat and salami containing pork. Serogroup O121 was the most prevalent non-O157 serotype found among pork carcasses. STEC O121 was previously linked with many outbreaks (4).

The potential of other strains isolated in our study to cause illness in humans should not be excluded. Serotypes O8 (1 EHEC and 25 STEC containing samples), O91(22 STEC containing samples), O139 (15 STEC containing samples), O20 (9 STEC containing samples) and O55 (7 STEC containing samples) were recovered. *E. coli* O8 possessing *stx*_*2e*_ has been reported to cause acute diarrhea (54), while O91 STEC strains can cause HUS or HC although they are *eae*-negative (55). In addition, O8, and O91 were included in the 20 most frequent serogroups reported in confirmed cases of human STEC infections in EU/EEA, 2015-2017 (3).

The results of the present study, observed with sampling only in two plants in the central part of the U.S. showed that pigs carry a variety of different STEC and EHEC serotypes, some of those serotypes are of high public health importance (e.g. O157 and O121), cross-contamination can occur during processing and dressing and interventions applied before chilling have an important role in reduction of microbial loads (APC, EBC) and prevalence of STEC and EHEC.

This study confirmed that market pigs presented for harvest in the U.S. carry a variety of STEC and EHEC serogroups, in decreasing order (O157, O121, O8, O91, O139, O20 and O55) and introduce them to the processing plant environment. Results showed that pork skin may be a significant source of EHEC and STEC to meat. The highest APC and EBC levels on pork skins were found during spring and summer, while the prevalence of genetic markers indicating the presence of STEC and EHEC were significantly less during winter. Hygienic processing at both plants significantly reduced contamination on carcasses, regardless of the season. Post-scald carcasses showed that STEC prevalence (indicated by presence of *stx* gene) was significantly decreased by 80-90% which makes this processing step key to contaminant reduction. Important control measures present included decontamination of pork carcasses with 2% lactic acid applied before chilling. Since the results from present study showed that higher prevalence of STEC and EHEC was detected during spring, summer and fall, a risk-based food safety management options should be implemented during these three seasons to achieve beneficial effect in reducing the pathogen prevalence on pork carcasses. Further in depth studies are needed to understand the sources of STEC and EHEC carried by pigs presented for harvest, cross-contamination of pork carcasses in the processing plant, and the impact of blast chilling on arresting the growth of bacterial contaminants on pork carcasses.

## MATERIALS AND METHODS

### Meat establishments

The sample collection was conducted in two establishments (Plant A and Plant B) approved for export of pork meat and deli-meat products to foreign markets by the USDA Food Safety and Inspection Service (FSIS). The selected meat establishments were two large US commercial hog processing plants that harvested 11000-17000 hogs/day. The harvest process and dressing operations followed standard procedures of: stunning, exsanguination, pre-scalding wash, scalding at 60 °C, dehairing, singeing, polishing, pre-evisceration wash, evisceration, carcass splitting, trimming, final wash, and chilling (final carcass and cooler temperature was 4 °C/16-24 h; Fig. 1). Plants A and B had different chilling systems, conventional and a blast chilling system, respectively.

**FIG. 1.**
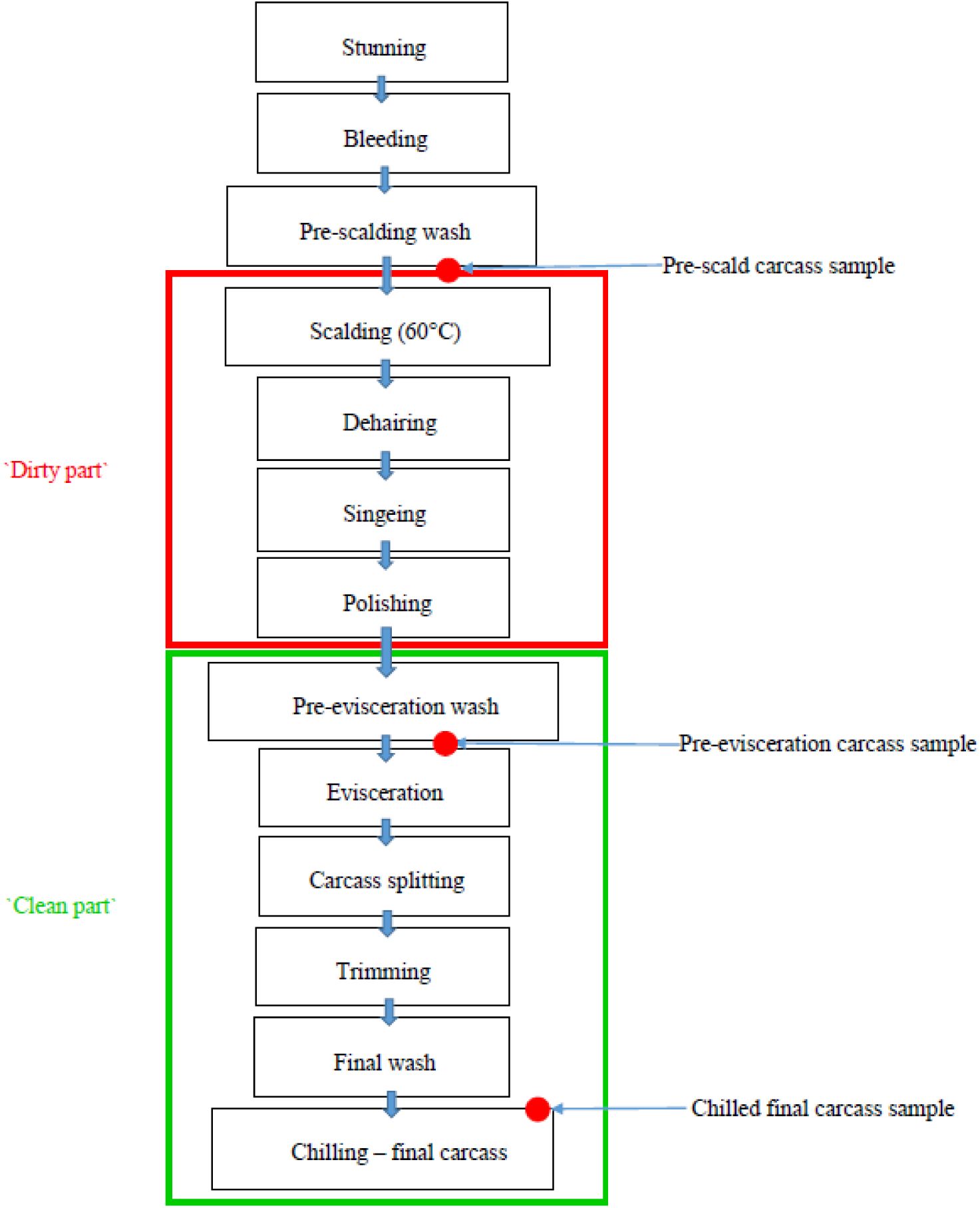
Pork slaughter line: standard operating procedures (SOPs) and sampling sites

### Sampling collection

The sampling protocol targeted the incoming contaminants on skins, then examined carcasses at two relevant locations: post-scald pre-intervention carcasses and finished carcasses after chilling. Therefore, identifying along the harvest line where pork carcasses may have been cross-contaminated with microbes, including STEC and EHEC (Fig. 1).

The sampling was carried out quarterly, throughout the year, covering four seasons, e.g. Q I - winter (December–February), Q II – spring (March–May), Q III - summer (Jun–August) and Q IV – fall (September–November). Each plant (designated Plant A and Plant B) was visited once per season and carcass samples were collected over two consecutive days on each trip, totaling eight sampling days per plant/per year, for a total of 16 sampling days/per year for two plants. On each sampling day, 95 samples were taken from three sampling points along the harvest line: skin of stunned exsanguinated pre-scald carcass, post-scald pre-evisceration carcass, and chilled final carcass. In total, 1536 samples were collected over the course of the study, 384 samples in each season (winter thru fall).

Samples were collected as described previously (using moistened cellulose sponges (Whirl Pak; Nasco, Fort Atkinson, WI), prewetted with 20 mL of buffered peptone water (BPW; Difco, Becton Dickinson, Franklin Lakes, NJ) (56). To prevent cross contamination, gloves were worn during sampling and were changed following each sample.

Samples from skin of pre-scald carcass surfaces were obtained by using both sides of the pre-wetted sponge to swab an area of approximately 1500 cm^2^ along the belly midline. After scalding, singeing, and polishing of the carcass, pre-evisceration post-scald carcass samples were obtained by using both sides of the pre-wetted sponge to swab approximately 4000 cm^2^ of the carcass surface along the midline from ham to sternum, including fore shank and jowl. Final carcass samples were obtained from carcasses that had been chilled at least overnight in coolers, by using both sides of the sponge to swab approximately 4000 cm^2^ of the carcass surface along the split midline from ham collar to jowl and fore shank. Due to the intense processing speed, in-plant operations and safety considerations for personnel collecting the samples, only a convenience sample was collected, therefore samples taken from each point were not matched to specific animals or groups of animals at other points. Skins and post-scald carcass samples were collected at the same time, while final carcass samples were collected after 24 h of chilling, from carcasses harvested on previous day. All samples were transported in coolers with ice packs (at <4 °C), received, and processed at the U.S. Meat Animal Research Center (Clay Center, NE, USA) within 24 h of collection, according to the protocol described by Schmidt et al. (56). The levels of Aerobic Plate Counts/APC (57) as hygiene level indicators, *Enterobacteriaceae*/EBC (58) as indicators of fecal contamination, and STEC non-O157 as pathogen (59) were determined.

### Sample processing

Each sponge swab was massaged by hand to ensure it was thoroughly mixed, then 1 mL was removed for APC and EBC. Eighty milliliters of tryptic soy broth (TSB; Difco, BD) was added to the remainder of the sample and sponge to enrich the samples for STEC. Enrichment consisted of incubation in a programmable incubator at 25 °C for 2 h, 42 °C for 6 h, then held at 4 °C until processed. After enrichment, two 1 mL portions of each sample were removed for STEC screening and analysis, with one of the portions archived as a frozen (−70 °C) 30% glycerol stock.

### Screening for Shiga toxin genes

One hundred microliters of an enrichment were placed in a microcentrifuge tube and used to prepare a crude DNA boil prep lysis (60). Two microliters of the DNA preparation were placed into separate 25 µL multiplex PCR reactions that detected *stx*_*1*_, *stx*_*2*_, *eae*, and *ehx* and was performed as previously described (61). Products of the PCR amplifications were separated by electrophoresis, stained using ethidium bromide, and then photographed and interpreted for the presence of the four possible reaction products. Enrichments that had *stx*_*1*_ and/or *stx*_*2*_ were considered positive for STEC, while enrichments that had *eae* and *stx*_*1*_ and/or *stx*_*2*_ were considered positive for EHEC for use in prevalence calculations.

### Isolation of STEC and EHEC

The sample enrichments determined by PCR to contain *stx*_*1*_ and/or *stx*_*2*_ were assayed by spiral plating of samples onto plates of washed sheep blood agar containing mitomycin (wSBAm) (62). Each enrichment was serially diluted to 1:500 and 1:5000 in cold (4 °C) buffered peptone water (BPW). Fifty microliters of each dilution were spiral plated, on to wSBAm plates. The plates were incubated overnight at 37 °C and then viewed on a white-light box for the suspect enterohemolytic phenotype as a thin zone (≤ 1 mm) surrounding the colony (63). In addition, if other hemolytic phenotypes such as alpha, beta, or gamma hemolysis were present, additional colonies representative of each hemolytic phenotype were picked for screening. A minimum of 4 colonies (if colonies were present) and a maximum of 6 colonies per sample were picked and placed into individual wells of 96-well screening plates containing 100 µL TSB per well. After suspect colonies were picked, the wSBAm plates were placed at 4 °C. The 96-well screening plate was incubated at 37 °C overnight, then screened by PCR as described above. If at least one suspect colony from a sample did not contain *stx1* and/or *stx2*, the wSBAm plates were removed from 4 °C and subjected to another round of suspect colony picking. All *stx*-containing isolates were checked for purity by streaking for isolation on sorbitol MacConkey agar containing 5-bromo-4-chloro-3-indolyl-#-D-glucuronide (SMAC-BCIG; Oxoid-CM0981; Remel Inc., Lenexa, KS) then transferred to tryptic soy agar (TSA; Difco, BD) plates for characterization.

### Characterization of isolates

All *stx*-containing isolates (STEC) and *stx*- and *eae*-containing isolates (EHEC) were confirmed to be *E. coli* by biochemical assays using Fluorocult LMX broth (Merck KGaA, Darmstadt, Germany) and API 20E strips (bioMerieux Inc., Hazelwood MO), both used according to the recommendations of the manufacturers. Once an isolate was established as being a STEC or EHEC, its serotype was determined by molecular and serologic identification of the O serogroup. PCR was used for molecular identification of O groups O26, O45, O55, O103, O111, O113, O117, O121, O126, O145, and O146 as described previously (64). *E. coli* antisera (Cedarlane Burlington, NC) were used to confirm the PCR results and identify other O serogroups. Virulence genes of each STEC or EHEC isolate were determined by PCR as described previously (64). Shiga toxin subtypes of the isolates were identified to be *stx*_*1a*_, *stx*_*1c*_, *stx*_*2a*_, *stx*_*2c*_, *stx*_*2d*_, and *stx*_*2e*_. If an *stx* subtype could not be identified simple the isolated was identified as “*stx*_*1*_” or “*stx*_*2*_”. Intimin (*eae*) subtypes: α1, α2, β1, β2, γ, δ, ε, θ, and ζ were identified by PCR as described previously (64) and if an *eae* subtype could not be identified for an isolate, it was referred to as ‘*eae*’. The presence of four genes associated with the large 60-MDa virulence plasmid, *toxB, espP, katP*, and *etpD*; additional toxin-encoding genes (*subA, lifA, cnf*, and *astA*); adherence-encoding genes (*iha* and *saa*); and hemolysin genes (*hylA* and *chuA*), were identified amongst the isolates by PCR as described previously (64). Lastly, genes described for molecular risk assessment associated with *E. coli* O157:H7 O-islands 36, 57, 71, and 122 (*nleB, nleE*, ent*G2-3, G5-2*, and *G6-2, nleC, H1-1, nleB2, nleG, nleG9, nleF, H1-2, nleA*, and *G2-1* were identified by PCR as described previously (64).

### Statistical analysis

Results from the enumeration (APC and *Enterobacteriaceae* count) of bacterial groups were analyzed for each sample type (skin, post-scald carcass, and final carcass) using analysis of variance with the GLM procedures of SAS. The model included main effects of season and plant. For significant main effects (*P* ≤ 0.05), least squares means separation was carried out with the PDIFF option (a pairwise t test). The data for enumerations were log transformed before the analysis of variance. Pairwise comparisons of frequencies were made using the PROC FREQ and Mantel-Haenszel chi-square analysis of SAS.

Sample enrichments were sorted according to serotype and screening PCR positive reaction pattern (*stx*_*1*_, *stx*_*2*_, and *eae*) and comparisons of prevalence were examined using a one-way analysis of variance (ANOVA) and the Bonferroni multiple-comparison posttest. Comparisons of median values of the data sets were made using the Kruskal-Wallis test for nonparametric data and Dunn’s multiple-comparison post test. For data sets with only two groups of values, comparisons were made using either a two-tailed unpaired t-test or the Mann-Whitney U test for nonparametric data. For cases when pair-wise differences were made, the DIFFER procedure of PEPI software (USD, Inc., Stone Mountain, GA) was used. In all cases significance being defined at a *P* value of ≤ 0.05.

## ACKNOWLEDGMENTS

This project was funded in part by The Pork Checkoff. We are grateful to the participating processors for allowing access for sample collection. We thank Michael Guerini for his scientific contributions. We thank Marilyn Bierman and Jody Gallagher for administrative assistance. Thanks to Greg Smith, Frank Reno, Julie Dyer, Bruce Jasch, and Lawnie Luedtke for technical assistance. USDA is an equal opportunity provider and employer. Names are necessary to report factually on available data; however, the USDA neither guarantees nor warrants the standard of the product, and the use of the name by USDA implies no approval of the product to the exclusion of others that may also be suitable.

